# The physiological adaptation for the “fore-mid” four-legged walking behavior of the pygmy mole cricket *Xya sichuanensis*

**DOI:** 10.1101/2020.03.22.002675

**Authors:** Yi Zhang, Shuying Wang, Zhu-Jun Feng, Tong-Xian Liu, Chengquan Cao

## Abstract

Animals have developed numerous specialized biological characteristics due to selective pressure from the environment. The pygmy mole cricket *Xya sichuanensis* has well-developed saltatorial hind legs for jumping and benefits for its survival, but these legs cannot be used for walking. Therefore, the typical tripedal gait used by most insects with six legs is not possible, and *X. sichuanensis* walks exclusively using its fore and mid legs. In this study, we describe a “fore-mid” walking pattern in *X. sichuanensis.* Further, we sought to deepen our understanding of the biological and physiological adaptations of this “four-legged” insect. We found the positions of tarsi points relative to the ground, integrated hind leg-abdomen structure, thickened ventral cuticle, and leg movements during walking to all show a unique biological adaptation. Of interest, *X. sichuanensis* was observed to demonstrate four-legged walking, underlining the general theme that insects have strong plasticity at both physiological and behavioral levels. We suggest that on an evolutionary timescale, *X. sichuanensis* has developed behavioral characteristics such as optimized jumping behavior and a unique walking pattern alongside specialized anatomical adaptations to enable survival in a competitive environment. This study could help explain biological and physiological adaptations for insects’ behaviors with important implications for the study of diversity in insect locomotion.

## 1. Introduction

Animals’ locomotion is often used as a method to avoid danger and to rapidly adapt to environmental changes. Walking and jumping are two types of locomotion behaviors for many insect species, and help respond to different types of threats (Burrows 2003; 2009; Burrows and Picker, 2010). Jumping, as a strategy to escape predators or to launch into flight, normally requires particularly specialized legs (typically the hind legs), and this specialization often affects patterns of walking (Usherwood and Runion 1970; Burns 1973; Burrows and Sutton, 2013).

Most adult insects have six fully functional legs adapted to walk, and a tripedal gait is most common (Hughes 1952; Wilson 1966; Dickinson et al., 2000). During this standard movement process, for each step, three legs touch the ground in a triangle shape to yield a stable stance. In a stride cycle, the middle leg on one side and the fore and hind legs on the other side are placed on the ground to form a triangle, and the other three legs are lifted and moved forward; after these lifted legs reach their position to form a new triangle, the first three legs begin to lift and move forward in a continuous symmetrical cycle (Wilson 1966; Cruse 1976; Full and Tu 1991). However, this pattern is sometimes varied under conditions such as running, and insects can also adjust their gait to cope with the loss of one or more legs (Grabowska et al.,2012).

Quadrupedalism (walking with four legs), is alternatively used in many animal species, especially in vertebrate animals, including mammals and reptiles (Full and Tu 1991; Dickinson et al., 2000). For many animals, during walking, the motion of legs on either side of the body alternates, or alternates between the front and back legs. “Four-legged” walking insects have also been observed, such as mantis (Mantodea) (Roeder 1937), water striders (Gerridae) (Dickinson 2003; Hu et al., 2003), and brush-footed butterflies (Nymphalidae) (Wolfe et al., 2011); one pair of legs (normally fore legs) of these insect species are often adapted for seizing, predation, or is simply reduced in size and not used for walking. In most Orthoptera species, the hind legs are saltatorial and possess a well-developed femur muscle adapted for jumping. Although their hind legs are specialized and are used both for jumping and walking, Orthoptera species such as locusts or grasshoppers use an alternating tripod gait (Usherwood and Runion 1970; Burns 1973). In most cases, these species use all six legs when walking (Wilson 1966; Burns 1973).

The pygmy mole crickets are a small species of Orthoptera (Burns 1973; Burrows and Sutton 2012). They normally live in banks by fresh water and have been used as an environmental indicator for dynamic river systems in Europe (Münsch et al., 2013). These insects exhibited many special behaviors based on their biological structures (Burrows and Picker 2010; Burrows and Sutton 2012).

The pygmy mole cricket species has short wings and mole-like fore legs that can be used to build burrows for nesting (Burrows and Picker 2010). Pygmy mole crickets also have a pair of well-developed saltatorial hind legs like some other Orthoptera species that can be used for jumping from land or even from water to avoid threats (Burrows and Picker 2010; Burrows and Sutton 2012). The legs of the pygmy mole cricket, especially the hind legs, have been described in detail in previous studies as they relate to jumping behaviors: many unique and specifically developed structures have been documented in the hind femur and tibia for jumping. However, according to our findings, the hind legs are too specialized to be used for walking (Burrows and Picker 2010). We assumed that these biological features necessitate that pygmy mole crickets walk on only four legs, not using the typical tripedal gait.

Four-legged insects and their patterns of movement have been studied in some detail (Burns 1973; Hu et al., 2003; Wolfe et al.; 2011; Grabowska et al.,2012). However, the pygmy mole cricket is different from typical “mid-hind legs” insects, instead walking with “fore-mid leg” motion. In this study, we sought to understand the biological adaptions of the pygmy mole cricket that allow four-legged motion and characterized its walking pattern, through biological parameters analysis, SEM scanning, HE staining and walking pattern analysis. This study could be helpful for understanding the biological adaptations that underlie insects’ behavior and carries new insights relevant to the study of walking in insects.

## 2. Materials and Methods

### 2.1. Insects

Pygmy mole crickets *Xya sichuanensis* (Cao et al, 2018) were collected from Leshan, Sichuan providence, China (29.5751° N 103.7534° E). The insects were kept in containers with a base of moist sand under long-day conditions (16L:8D; 20 ± 1°C; > 80% RH) at the College of Life Science, Leshan Normal University, Leshan, Sichuan, China, and experiments were conducted at the Key Laboratory of Applied Entomology, Northwest A&F University, Yangling, Shaanxi, China.

### 2.2. Images collection

Digital image acquisition and body length measurement were performed using a Panasonic DMC-GH4 digital camera (Panasonic, Osaka, Japan) attached to a dissecting microscope SDPTOP-SZN71 system (Sunny, Hangzhou, Zhejiang, China). The parameters were measured by ImageJ software (Wayne Rasband, National Institutes of Health, Bethesda, Maryland, USA)

### 2.3. SEM and Histological sectioning and staining

The anatomy of the legs and abdomen were examined by electron microscopy and histological sectioning and staining. Samples for scanning electron microscopy (SEM) were treated with 2.5% glutaraldehyde fixative for 24 h, ultrasonically cleaned for 3 min, and washed with 3 mol/L phosphate buffer (pH = 7) 3 times (10 min/time). Next, samples were placed in dehydrated stepwise using ethanol (70%, 80%, 90% and 100%) and, finally, dried at room temperature for 12 h. After angle adjustment, the samples were sputter-coated with gold and images were taken under scanning electron microscope (Accelerating voltage: 10.0 kV).

Histological sectioning and staining were performed at the third abdominal segment (A3) of adult *X. sichuanensis* in cross section. The samples were fixed in 10% (v/v) buffered formalin overnight, dehydrated, embedded in paraffin, and sectioned. Slides were prepared by soaking in xylene twice for 20 min, 100% alcohol twice for 5 min, and 75% alcohol for 5 min, followed by rinsing with water. The slides were then immersed in hematoxylin solution for 3–5 min and rinsed with water. The sections were destained with acid alcohol and rinsed, treated with ammonia solution and rinsed in slowly-running tap water, and then placed in 85% alcohol for 5 min, 95% alcohol for 5 min, and eosin for 5 min. The sections were dehydrated 3 times with 100% alcohol (each for 5 min), treated with xylene twice (each for 5 min), and mounted with resin. Digital images were acquired using a Nikon DS-Ri1 camera (Nikon, Tokyo, Japan), a Nikon 80i microscope system (Nikon, Tokyo, Japan), and Nis-Elements v. 3.22.14 (Build 736, Nikon, Tokyo, Japan). Cuticle thickness was determined from the digital images. Four different parts of the abdominal cuticle were measured. Abdominal sections of wingless *A. pisum* and *B. germanica* (both of them are “walkers” and cannot fly) were also performed for cuticular thickness detection. Cuticular thickness were calculated by software Nis-Elements v. 3.22.14.

### 2.4. Walking patterns

Walking patterns of *X. sichuanensis* were analyzed from video recordings. In order to prevent jumping during video collection, the hind legs of *X. sichuanensis* were removed, and then insects were allowed to recover for 48 hours before experiments. Video files were recorded for ~5 min using a Panasonic DMC-GH4 digital camera (Panasonic, Osaka, Japan) with a macro lens (Canon^®^ Macro lens EF 100 mm 1:2.8 L IS USM, Canon, Japan; equivalent focal length: 200 mm). The camera was set to high-speed recording (96 fps) at the highest resolution (1920X1080). Movement speed and leg movements were analyzed from recorded files by ImageJ and EthoVision XT (Noldus Information Technology, Wageningen, the Netherlands).

### 2.5. Statistical analyses

Values of biological parameters were subjected to one-way analysis of variance (ANOVA). Differences among means were calculated using Duncan’s test at a significance level of *P* < 0.05.

## 3. Results

### 3.1. Legs

We found *X. sichuanensis* fore legs to be the shortest (femur, 0.722 ± 0.013 mm; tibia, 0.529 ± 0.011 mm; N = 5, Fig. 1C), and the line connecting two fore tarsi points intersected the midline inside the body (Fig. 1G).

**Fig. 1.**
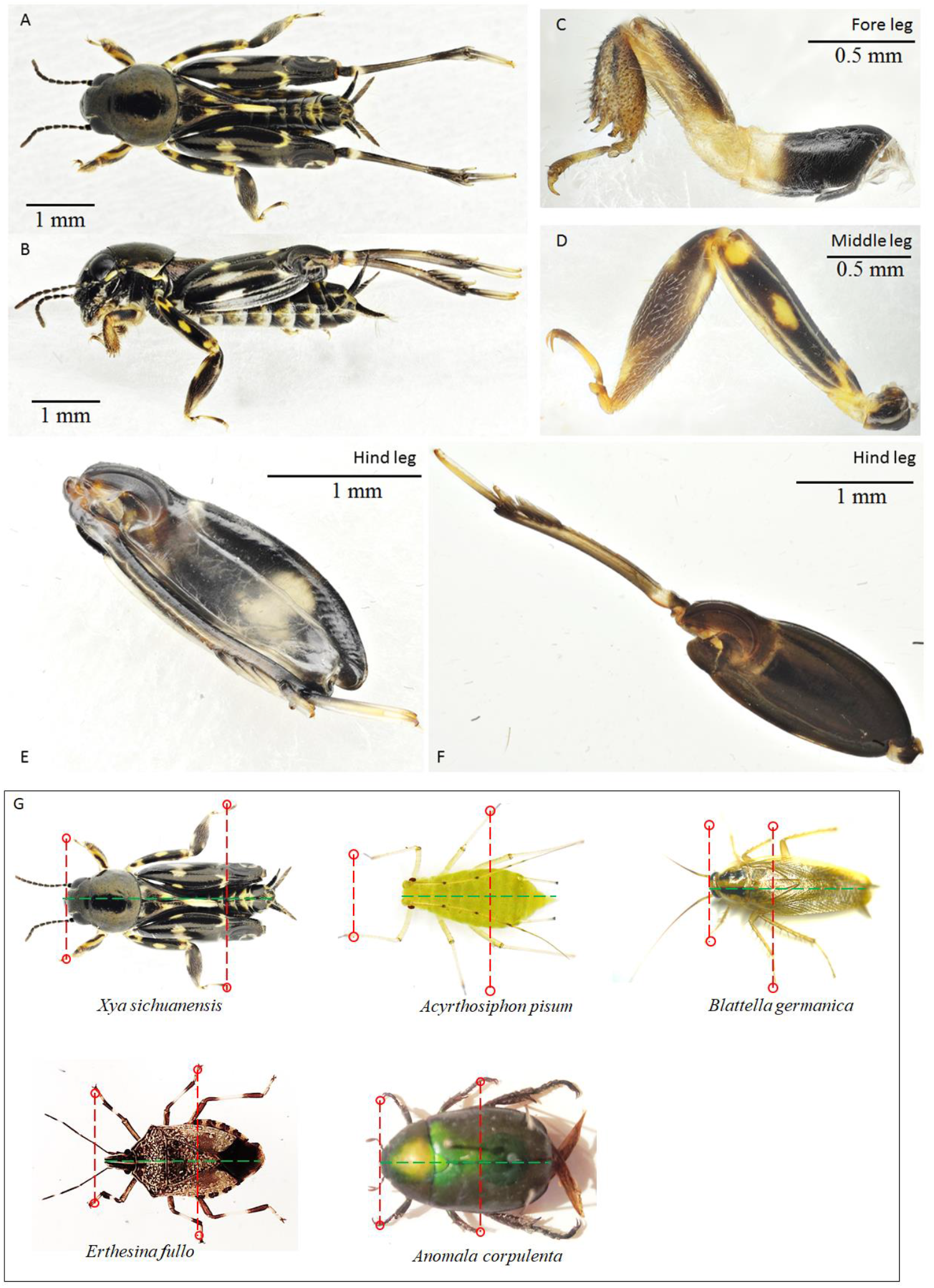
Body structure of the pygmy mole cricket *Xya sichuanensis.* Dorsal and side view of whole body are shown in A and B. Magnified legs are shown in C, D, E and F. The length ratio of two parts to intersection of the midline and the line connecting of two middle tarsi (anterior part/posterior part; 4.037) of *X. sichuanensis* and other selected insects *(Acyrthosiphon pisum, Blattella germanica, Erthesina fullo, Anomala corpulenta,* reared at the Key Laboratory of Applied Entomology or collected from University Museum Garden, Northwest A&F University, Yangling, Shaanxi, China) are shown in G.

The middle legs are relatively longer (femur, 1.318 ± 0.030 mm; tibia, 1.139 ± 0.043 mm, N = 5, Fig. 1D) and their tarsi are located posteriorly. The length ratio of two parts to intersection of the midline and the line connecting of two middle tarsi (anterior part/posterior part; 4.037) of *X. sichuanensis* is notably higher than other selected insects, underlining the comparatively wider supporting area for four legs of *X. sichuanensis* relative to other insects *(Acyrthosiphon pisum,* 1.226; *Blattella germanica,* 0.947; *Erthesina fullo,* 1.409; *Anomala corpulenta,* 1.483, Fig. 1G).

The un-flexed hind legs of *X. sichuanensis* were found to be longest, with measurements of 2.292 ± 0.026 mm for the femur and 1.869 ± 0.478 mm for the tibia (N = 5). The flexed hind legs exhibited no contact with the ground in either stationary or moving states (Fig 1E & F).

The tibia fits in a groove of the femur when the hind leg is fully flexed, and three cover plates of each sides of the femur are cover the pivot joint (Fig. 2A & B). Meanwhile, we noted that the entire flexed hind legs could also be grooved along the dorsal surface of the abdominal segments (Fig. 2C). The triangular wings (both fore and hind wings) of *X. sichuanensis* are short (2.09 ±0.055 mm in length) and cover less than 2/3 abdominal segments by length (3.57 ± 0.037 mm, without cerci, Fig. 2D-F). All of these parts fit together to form an integrated structure (Fig. 2C & E).

**Fig. 2.**
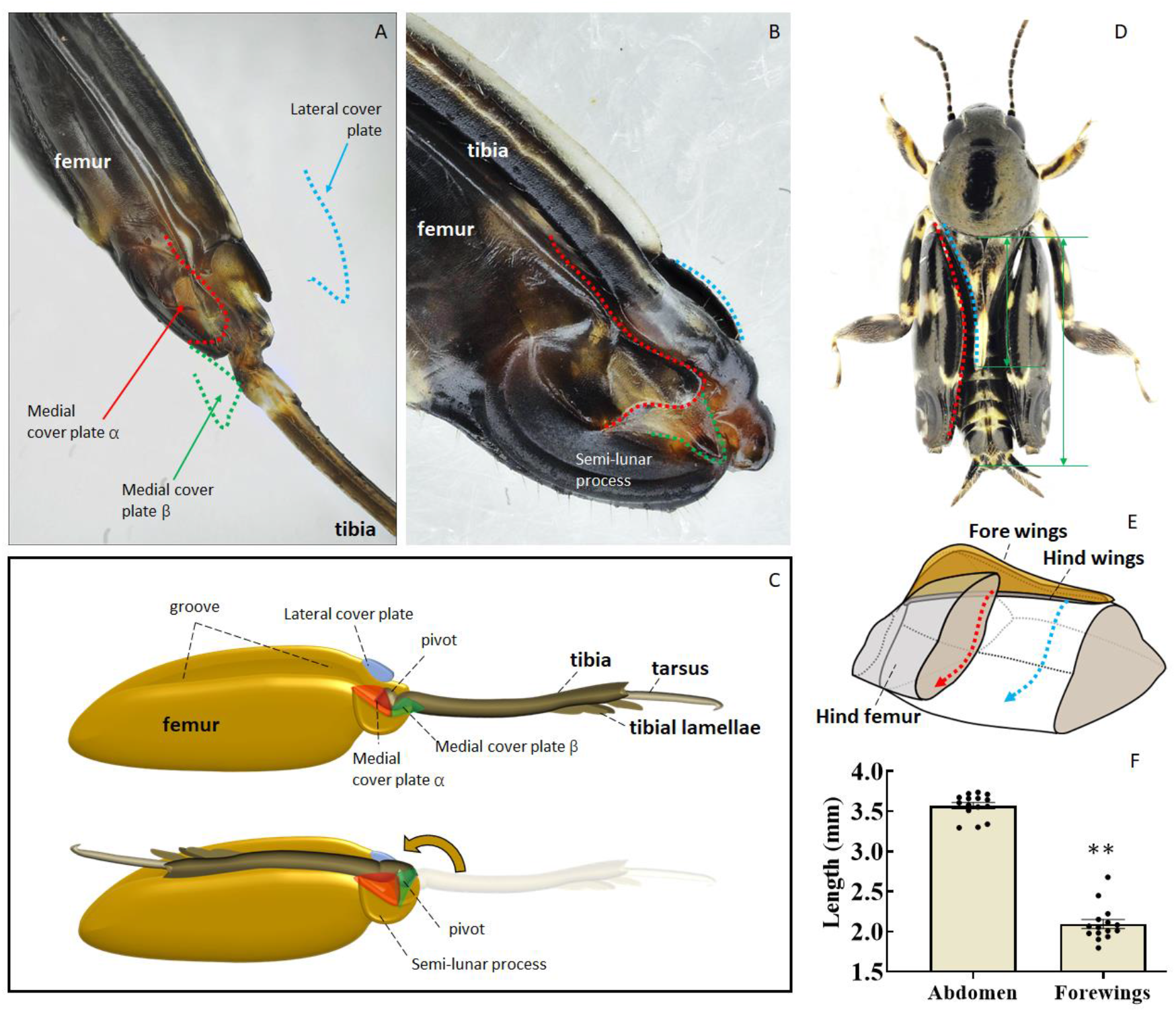
Images and sketches of the structures of the hind leg. The femoro-tibial joint of a hind leg is shown in A and B. A sketch of a hind leg is shown in C. Length ratio and structural sketches of the hind leg and abdomen are shown in D, E and F. ** in panel F indicates significant difference at *P* < 0.001 (Student’s *t*-test).

### 3.2. Abdomen

The abdominal ventral cuticle segments exhibit flat surfaces (especially A2, A3 and A4 segments) and make contact with the ground (Fig. 3A & B, could also be found in Fig. 4B). The hind leg coxas of both sides show a similar flat surface (Fig. 3B). Two types of trichome structures can be seen in the abdominal ventral cuticle; the longer trichome structure is about 183.7 ± 9.5 μm, while the short one is about 66.6 ± 1.8 μm. The scale-like cuticle units with small convex could be detected under high magnification (Fig. 3C & D).

**Fig. 3.**
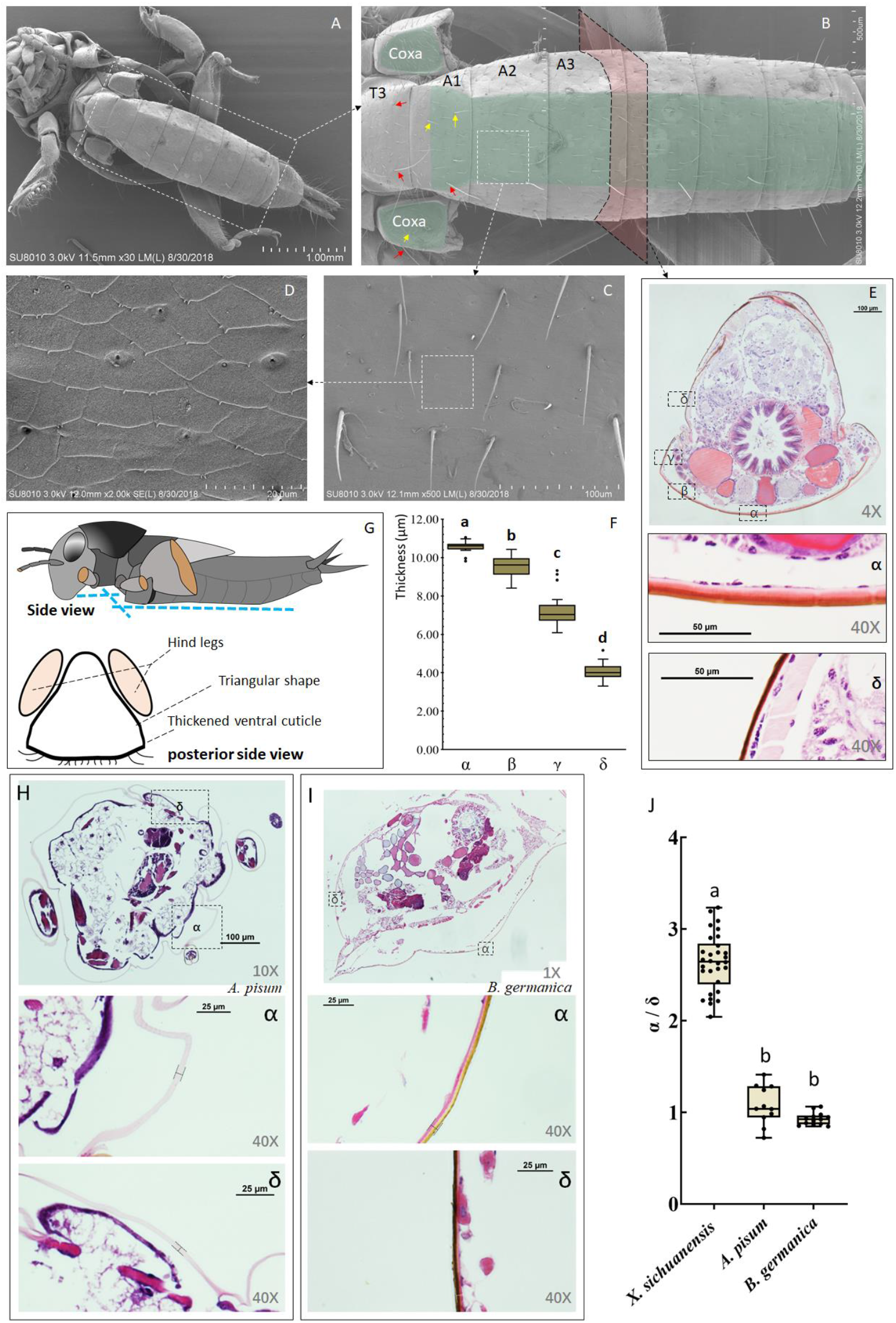
Scanning electron micrograph (A-D) and histological sectioning of abdominal cross section (E) of the abdomen. The thickness of the abdominal ventral cuticle is shown in E-α and E-β, and quantified results are shown in F; different letters above bars in panel F indicate significant differences in values (ANOVA, Duncan’s test, *P* < 0.05). Sketch of abdomen from the side and posterior side views are shown in G. Histological sectioning of abdominal cross section of *A. pisum* (H) and *B. germanica* (I). The thickness ratio of ventral abdominal cuticle (α) and side abdominal cuticle (δ) among three insect species *(X. sichuanensis*, *A. pisum* and *B. germanica*) shows in J.

Histological sectioning of abdominal crosscutting showed a varied thickness of the cuticle at different positions. The cuticle at the ventral position (which is in contact with the ground and measured about 10.6 μm on average) is significantly thicker than those of other locations *(F* = 715.834; df = 3, 116; *P* < 0.001; Fig. 3E & F). Meanwhile, compared with *X. sichuanensis*, the thickness differences between ventral cuticle and side cuticle were significantly lower in some other insect species (*A. pisum* and *B. germanica; F* = 281.09; df = 2, 51; *P* < 0.001; Fig. 3H, I&J).

### 3.3. Walking

We observed two types of walking patterns in our studies: fast-walking (running) and slow-walking. The average speed of fast-walking was approximately ten times faster than slow-walking *(T* = −21.801; df = 33; *P* < 0.001; Fig. 4A), and the associated patterns of leg movement were also notably different.

#### 3.3.1. Fast-walking (running)

The fast-walking process of *X. sichuanensis* was divided into four steps. In a stride cycle, it started with movement of the left fore leg (L1). The right middle leg (R2) was lifted before L1 reached its target point ahead; right fore leg (R1) and left middle leg (L2) are pivot legs at this stage. Meanwhile, the abdomen was involved as a fulcrum point. Similarly, after L1 and R2 reached their target points, R1 and L2 mirrored that same movement (Fig. 4C). The alternation of four steps in one cycle continued during straight running (Fig. 4D&E). The movements of the left fore leg and right mid leg occurred at the same time during one phase, and the right fore leg and left mid leg moved at the same time in the second phase (Fig. 4F).

**Fig. 4.**
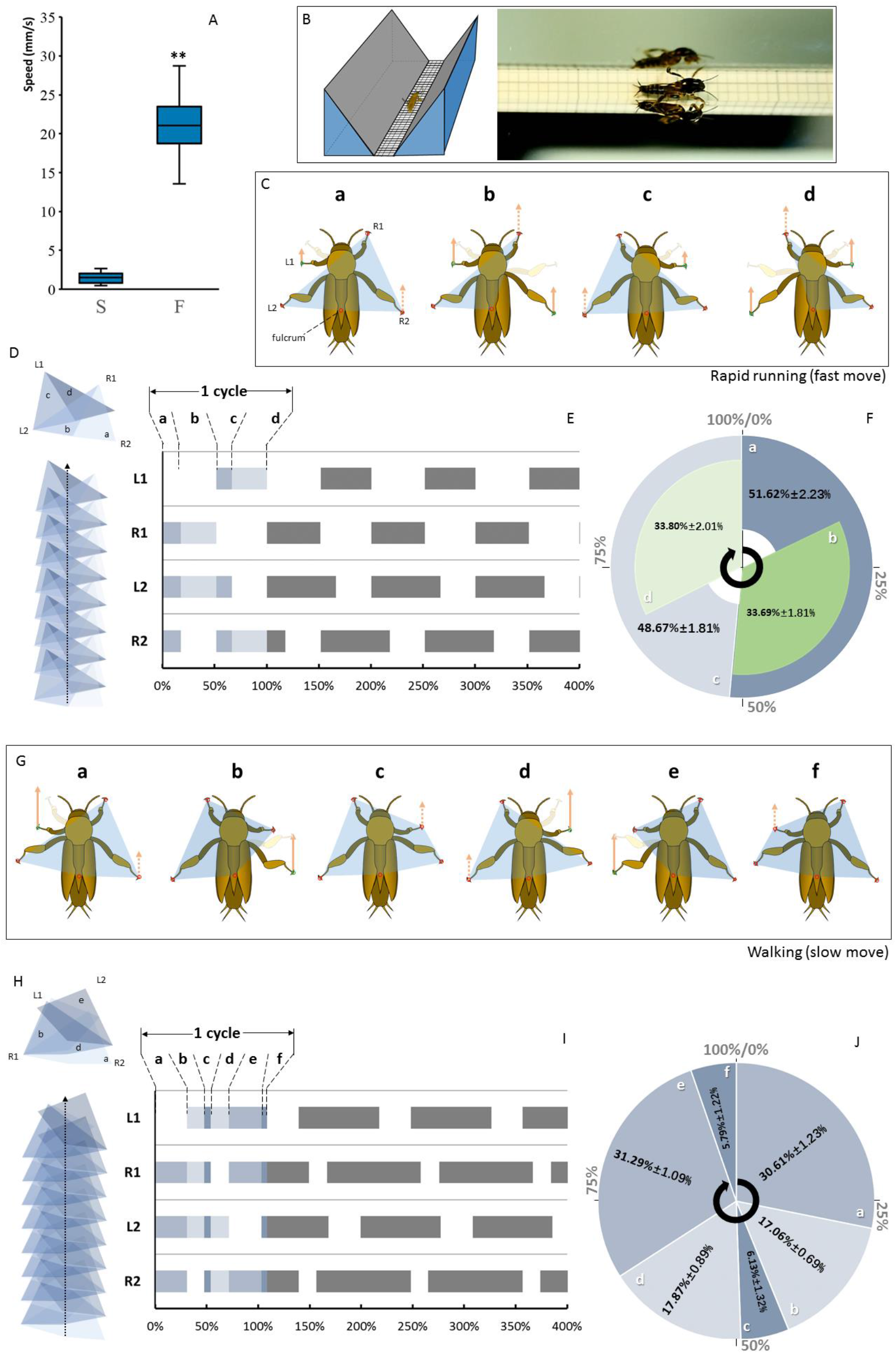
Two types of walking patterns of *Xya sichuanensis* were measured in our data. The observation device used for these studies is shown in B. Fast-walking is shown in C-F, and slow-walking is shown in G-J. The individual walking stages for the two types of walking are shown in C and G, the ground-contact polygons a marked in the images and then merged and overlaid to create moving sequences (D and H); alternating stepping patterns during two types of walking are shown in E and F, and proportions of each step’s time spent in each moving cycle were calculated and shown in F and J. Lower-case letters in D-F (a, b, c and d) represent corresponding stages shown in C, and lower-case letters in H-J (a, b, c, d, e and f) represent corresponding stages shown in G. The velocity difference between these two types of walking is shown in A; ** indicates significant difference at *P* < 0.001 (Student’s *t*-test).

#### 3.3.2. Slow-walking

The slow-walking process of *X. sichuanensis* was divided into six steps. In a stride cycle, it also started with L1, however R2 was not lifted until L1 reached its target point. After R2 reached its target point, there was a short delay before R1 was lifted during which all five supporting points (four tarsi and abdominal ventral cuticle) were in contact with the ground; Similarly, in the next stage, R1 was then lifted, and L2 did not lift until R1 reached its target point (Fig. 4G-J).

## 4. Discussion

In this study, we investigated the biological features and physiological adaptations underlying the unique walking movements of the pygmy mole cricket *X. sichuanensis.* The positions of tarsi points with respect to the ground, the integrated hind leg-abdomen structure, the thickened abdominal ventral cuticle, and the pattern of leg movement during walking all show a specific pattern of development and biological adaptation to allow four-legged walking in the context of specialized jumping.

*Xya sichuanensis* is not the only insect species walked by four legs, however, it is not a typical four-legged” insect species; in contrast with other “four-legged” insects (mid-hind legged walking; Roeder 1937; Dickinson 2003; Hu et al., 2003; Wolfe et al.; 2011), *X. sichuanensis* uses its fore and mid legs for movement, and the abdomen is also used to support the body. Together, there are five supporting anatomical points during walking. *X. sichuanensis’s* fore legs are reported to be developed for digging and burrowing, and its mid legs are adapted to contact ground posteriorly for better walking using only four legs. It is clear that the great extension of the four legs (especially the mid legs) and ground-touching abdomen could offer a relatively wider supporting area and better stability during “four – legged” walking. This finding preliminary explained the feasibility that *X. sichuanensis* could use only four legs for its supporting or even walking. Similar phenomenon could also be observed in some other insect species, which their fore and mid legs have a greater participation at body supporting and walking (Usherwood and Runion 1970; Burns 1973).

*Xya sichuanensis* has six fully functional legs, compared with fore and mid legs, we found that the hind legs of *X. sichuanensis* exhibited no effect on walking and are mostly flexed on the dorsal side during walking. The function of the hind leg appears to be relatively independent and specialized for jumping like other Orthoptera species (Burrows and Picker 2010; Burrows and Sutton 2012). The uniquely developed structures of the legs and abdomen support this hypothesis. The short fore and hind wings of *X. sichuanensis* only cover part of the abdomen. This allows the whole flexed hind legs to also be grooved along the dorsal surface of the abdomen and fit together to form an integrated structure. We believe that there is a direct relationship between these specialized hind legs and specific morphological adaptations of the abdominal structure. Together, these studies reveal that the hind legs of *X. sichuanensis* have developed in support of optimized jumping and cannot be used for walking. The jumping behavior of *X. sichuanensis* possibly drives certain organs specialization and determines the developing direction. In this case, the importance of improved jumping outweighs six-legged walking, and indicating that jumping has a greater significance for its survival; and in turn, the special role of hind legs drives other four legs developing in support of walking.

The abdominal physiological characters of *X. sichuanensis* were also observed to possess special adaptations to allow four-legged walking. As discussed, the hind legs and abdomen fit together and the wings that cover the abdomen are shortened. Further, we noticed other special features in the ventral cuticle of abdominal segments (flat surfaces). Similarly, it was also observed at the ventral cuticle and the bottom of the hind coxas. We believe that, as this insect species has a relatively long cylindrical body shape, these specific angle variations among segments could support the abdomen’s use as a fifth fulcrum during walking. In this case, the ventral cuticular tissues of those structures is specialized for the function: a significantly thickened ventral cuticle observed, which was most likely adapted to protect the abdomen from friction during walking; and we did not detect this ventral cuticular thickening in other experimental species. It is a reasonable physiological character for special walking behavior of *X. sichuanensis*, and this specialization is believed to be one of the biological foundations for “abdominal supporting” during its working. Meanwhile, there were also two types of trichome structures detected at the abdominal ventral cuticle, which are possibly used for sensation during walking. It is obviously that the abdomen tissues have specialized in both morphology and functions to adapt to “four-legged” walking. These observed features from walking behavior to biological characteristics reflect an integrated structural and physiological adaptation to selective pressure from the environment.

Based on those biological features mentioned above, *X. sichuanensis* is well-suited for four-legged walking and very poorly suited for a typical tripedal gait (Hughes 1952; Wilson 1966; Dickinson et al., 2000; Goldman et al., 2006). We intent to figure out the details of *X. sichuanensis* walking with its four legs. Unfortunately, due to limited equipment capacity, only two types of walking patterns were identified in our data, and it should be varied for *X. sichuanensis* to adapt to complex walking conditions. These moving patterns were similar to the grasshopper *Tropidopola cylindrica*, which has also been observed to use the abdomen as a supporting fulcrum for walking (Wilson 1966). Although some other four-legged walking insect species, such as mantis generally uses its mid and hind legs for walking, the gait patterns between *X. sichuanensis* and mantis were relatively similar (Roeder 1937). It is a convergence adaptation on behavior level among different species; and these insect species happened to select this effective pattern to handle walking problem without another two legs (although the pairs of un-participated legs are different). The movement process and many biological features of *X. sichuanensis* appear to be perfectly suited to this “four-legged” walking. This phenomenon reveals the adaptability of a highly specialized behavior in support of survival. Combining with the importance of jumping behavior of *X. sichuanensis* that reported (Burrows and Picker 2010; Burrows and Sutton 2012), we believe that the emergence of this specialized walking behavior is fundamentally induced by environmental pressure that has increased the weight of jumping behavior for this species.

In brief, this study reveals that insects have strong plasticity at both the physiological and behavioral levels, which is expressed as a specialization of body structures, forms, and behaviors. We believe that characteristics that favor the survival of species will be preferentially preserved, such as the jumping behavior of *X. sichuanensis*, and that other biological characteristics will adapt in support of those specialized structures. We expect that similar characteristics may be found in other jumping insects. This finding could be helpful for understanding the biological adaptations underlying insect behavior, and provides new data to describe the diversity of walking among insects.

## Acknowledgments

We are grateful for the assistance of all staff and students in the Key Laboratory of Applied Entomology, Northwest A&F University, and students in College of Plant Protection, Shandong Agricultural University, China.

## Formatting of funding sources

Funding of this research was supported by the Chinese Universities Scientific Fund (grant number, Z109021718).

